# Single-Cell and Spatial Transcriptomics Integration Identifies Mural Cell Oxidative Stress Genes *Clu* and *Gria2* as Key Biomarkers in Ischemic Stroke

**DOI:** 10.64898/2026.02.09.704987

**Authors:** Jiali Xie, Jiaxing Zhu, Chunyan Zhou, Jing Yao

## Abstract

Oxidative stress (OS) is a key factor in ischemic stroke (IS), but the characterization of OS-related genes in IS remains largely unexplored. Identifying key OS-associated genes could improve our understanding of OS in IS. We analysed single-cell RNA sequencing datasets and utilized AUCell, Ucell, singscore, ssgsea, and AddModuleScore algorithms, along with correlation analysis, identified 167 OS-related genes potentially linked to IS. Furthermore, we used seven machine learning algorithms, including least absolute shrinkage and selection operator, XGBoost, Boruta, random forest, gradient boosting machines, decision trees, and support vector machine recursive feature elimination, to identify the optimal feature genes: Clusterin (*Clu*) and Glutamate Ionotropic Receptor AMPA Type Subunit 2 (*Gria2*). Bulk RNA-sequencing showed that *Clu* expression was upregulated and *Gria2* expression was downregulated in IS tissues. Single-cell analysis further revealed that these expression changes predominantly occurred in mural cells, emphasizing their cell-specific roles in IS. Subsequently, we validated these findings through spatial transcriptomics (ST) analysis and animal model experiments. These results suggest that *Clu* and *Gria2* may participate in the progression of IS by altering the OS activity of mural cells. However, further research is needed to validate its clinical efficacy and ensure its application in the treatment of IS.

## 1. Introduction

Ischemic stroke (IS) is an acute cerebrovascular disease caused by occlusion of cerebral blood vessels, often leading to symptoms related to neurological dysfunction, including motor, sensory, visual, speech, and cognitive impairment. As the second leading cause of death and the primary cause of disability globally, IS poses a critical public health challenge due to its high incidence, disability, and mortality rates. The essence of this disease is ischemic and hypoxic damage to brain tissue caused by hemodynamic disorders, which then triggers complex cascade pathophysiological reactions through pathological mechanisms such as free radical damage, excitotoxicity, and inflammatory cascade reactions (1). Oxidative stress (OS), as a pivotal aspect in the pathological process of IS, has garnered considerable attention in recent years (2). OS refers to the imbalance between the oxidative and antioxidant systems in the body, leading to excessive production of reactive oxygen species (ROS) and reactive nitrogen species, which in turn damages biomolecules such as lipids, proteins, and nucleic acids in cells (3). During the occurrence of IS, the ischemic and hypoxic state of brain tissue can induce mitochondrial dysfunction, NADPH oxidase activation, and enhanced xanthine oxidase pathway, among other mechanisms, leading to the massive generation of ROS. These excessive ROS not only directly attack the polyunsaturated fatty acids in cell membranes, triggering lipid peroxidation reactions that compromise membrane integrity and fluidity, but also induce oxidative reactions with proteins, leading to structural alterations and functional inactivation. Consequently, these changes disrupt intracellular signaling transduction and metabolic regulation, thereby impairing essential physiological processes (4, 5). Moreover, DNA, as the genetic material within cells, is also highly susceptible to oxidative stress-induced damage. Oxidative damage to DNA can result in a variety of mutations and strand breaks, thereby disrupting normal gene expression and cellular proliferation and differentiation. Ultimately, these alterations may lead to apoptosis and necrosis of the cells (6). OS plays a crucial role throughout the entire course of IS, from the early injury of brain tissue during ischemia to the further damage following reperfusion. Investigating the mechanisms of OS-related genes in IS not only aids in a comprehensive understanding of the pathophysiological processes involved but also offers novel insights and potential targets for the development of targeted therapeutic strategies.

At present, single-cell RNA sequencing (scRNA-seq) has been widely used to analyze cell heterogeneity and reveal gene expression differences under different cell types and states (7, 8). Spatial transcriptomics (ST), as a newly established method, enables the precise localization of gene expression at the tissue level, providing spatial context to the transcriptome (9). The combination of scRNA-seq and ST has the advantage of accurately annotating specific cell subpopulations and revealing their spatial characteristics (10). Machine learning significantly enhances the advantages of scRNA-seq analysis by optimizing the analysis process of complex data and deeply integrating with bioinformatics methods. It not only enhances the ability to analyze datasets, but also assists in identifying key regulatory genes and pathways, thereby improving the accuracy of data analysis and revealing hidden relationships in the data (11, 12). In this study, we integrated scRNA-seq and bulk RNA sequencing (Bulk RNA-seq) to investigate genes related to OS in IS. Through machine learning-driven feature selection genes associated with OS, followed by validation through ST analysis and animal model experiment, thereby confirming the robustness of these findings and their pathophysiological relevance. This study provides new insights into the mechanism of OS-related genes in IS and advances the understanding of their potential molecular mechanisms. These findings provide a significant theoretical basis for novel therapeutic strategies of IS.

## 2. Methods

### 2.1. Data acquisition

The data source and screening process for this study are as follows: scRNA-seq data was obtained from the Gene Expression Omnibus (GEO) database GSE174574 dataset (https://www.ncbi.nlm.nih.gov/geo/), which includes 3 middle-cerebral artery occlusion (MCAO) treated and 3 sham operated mouse brain tissue samples. The Bulk RNA-seq data used the GSE233811 dataset as the discovery queue, which includes RNA expression profiles of brain tissues from 28 MCAO treated and 20 sham operated mouse. And select the GSE137482 dataset to construct an external validation queue, including 12 MCAO treated and 12 sham operated mouse brain tissue samples. The spatial transcriptome data was collected using the GSE233813 dataset.

Based on the GeneCards database (v5.24) - an integrated bioinformatics platform that integrates authoritative biomedical knowledge bases such as OMIM, PubMed, and Gene Ontology (GO), according to previous research (13), 2054 high confidence OS related genes were screened by setting the OS-related score threshold to ≥7 (Table S1). This screening criterion ensures that candidate genes have clear biological associations, laying the foundation for subsequent mechanism research.

### 2.2. scRNA-seq dataset analysis

In this study, the Seurat package (version 5.0) was used for data integration and standardization (14). The raw scRNA-seq data was subjected to quality control using the following criteria: The percentage of mitochondrial genes (pctMT) is less than 20%, the number of detected genes is between 200 and 8000 per cell, and the red blood cells gene expression level less than 3%. This process retained 58440 high-quality cells for downstream analysis. After logarithmic normalization, cells identify the top 2000 highly variable genes using the “FindVariableFeatures” function. Correct batch effects between samples through Harmony. Dimensionality reduction is achieved through principal component analysis (PCA), retaining the first 30 principal components for subsequent analysis. Use “FindClusters” for cell Clustering. Visualize through Uniform Manifold Approximation and Projection (UMAP). We annotated cell types using typical marker genes combined with CellMarker 2.0 (Table S2).

### 2.3. Evaluation of OS activity and enrichment analysis

This study used five single-cell scoring algorithm: AUCell (15) (UCell (16)) singscore (17)(ssgsea (18) and AddModuleScore (19) to quantify OS activity at the single-cell level and obtained a comprehensive activity score (20). Using the median score as the threshold for binary differentiation, we divided the cell population into high OS (oxidative stress) activity and low OS activity groups. We then identified genes significantly associated with OS activity through Pearson correlation analysis. Differentially expressed genes (DEGs) involving OS was performed using the “FindMarkers” function. By integrating the intersection genes from both DEGs and the correlation analysis, we generated a gene set for subsequent functional validation studies. Furthermore, we conducted GO enrichment analysis using the “ClusterProfiler” package to evaluate the potential roles of OS-related genes in biological processes, molecular functions, and cellular components (21).

### 2.4. Machine learning algorithms further identify the optimal feature genes

To identify the optimal feature genes most associated with OS activity, we used seven machine learning algorithms for feature selection and optimal gene recognition, including eXtreme Gradient Boosting (XGBoost) (22), random forest (23), Boruta (24), support vector machine recursive feature elimination (SVM-RFE) (25), gradient boosting machines (GBM) (26), decision trees (27), and least absolute shrinkage and selection operator (LASSO)(28). These algorithms ensure comprehensive and reliable identification of genes most related to OS activity through different feature selection strategies. Among them, LASSO enhances the accuracy and interpretability of the model through regularization and cross validation. Random forest utilizes random sampling and feature selection to construct multiple decision trees. Boruta processes high-dimensional data by comparing the importance of raw features with randomized features. XGBoost improves accuracy through iterative training and early stopping strategies; SVM-RFE enhances the interpretability and robustness of the model by recursively eliminating irrelevant features. GBM combines multiple weak learners to construct a powerful prediction model; The decision tree divides feature by maximizing subset homogeneity. By integrating these complementary algorithms, we have identified key central genes that contribute to OS activity with high confidence and robustness, providing a solid foundation for subsequent research and potential clinical applications.

### 2.5. The expression and predictive significance of the optimal feature genes in IS tissues

The expression analysis of the optimal feature genes was performed in two Bulk RNA-seq datasets (GSE57338 and GSE116250) using Wilcoxon rank sum test. To evaluate diagnostic performance, receiver operating characteristic (ROC) analysis was performed using the “pROC” package.

### 2.6. Validation of optimal feature genes at the single-cell level

By analyzing annotated scRNA-seq data, we explored the enrichment of optimal feature genes in various cell types. Through this analysis, we identified the specific cell types with the optimal feature genes involved in regulating OS activity. This method enables us to determine the cellular environment in which these genes function, thereby gaining a deeper understanding of their roles in the biological processes being studied.

### 2.7. Gene set enrichment analysis of the optimal feature genes

We conducted gene set variation analysis (GSVA) on the optimal feature genes using the GSVA package (version 2.1) to evaluate their correlation with potential specific biological processes or disease states, as well as potential regulatory mechanisms. GSVA is a non-parametric, unsupervised method that estimates changes in gene set enrichment in a sample by converting gene expression data into a gene set sample matrix. This method allows for the evaluation of pathway enrichment for each sample and facilitates further pathway centered analysis, such as differential expression, survival analysis, clustering, and correlation analysis.

### 2.8. Trajectory analysis

To infer single-cell trajectories and reveal cell state transitions, we used the Monocle package (version 2.3.6). We divide cells into two categories based on the presence or absence of optimal gene expression to construct cell trajectories (29). Monocle2 employs reverse graph embedding technology to reconstruct single-cell trajectories, thereby enabling robust and accurate analysis of complex biological processes. Using the UMI count matrices and the negbinomial.size parameter, we constructed a CellDataSet object under the default settings.

### 2.9. Cell-cell communication analysis

To explore the changes in intercellular communication modules, we integrated gene expression data using the CellChat package and used the default CellChatDB as the ligand receptor database (30). By identifying overexpressed ligands or receptors in each cell population and detecting enhanced ligand receptor interactions when ligands or receptors are overexpressed, cell type specific interactions can be inferred.

### 2.10. Spatial transcriptome analysis

We analyzed spatially resolved transcriptome data using the Seurat package (version 5.0). Exclude genes detected in 3 or fewer spots and standardize the count using SCTransform. Use the “FindTransferAnchors” and “TransferData” functions in the Seurat package to project single-cell RNA seq data onto spatially resolved data using default settings.

### 2.11. Animal model

The animal procedures in this study were approved by the Ethics Committee of Gannan Health Vocational College (Approval Number: 25001) and conducted in strict accordance with the Animal Research: Reporting of In Vivo Experiments (ARRIVE) guidelines and the National Institutes of Health (NIH) guide for the care and use of laboratory animals. We adhered to the 3R principles to minimize the number of animals used. Adult male Sprague-Dawley rats (60-80 days old, weighing 250-300 g) were purchased from Hunan Jingda Experimental Animal Center of China (license number: SCXK 2016-0002) and were subjected to a 12-hour light-dark cycle and had ad libitum access to standard rodent diet and water. Six rats were randomly divided into two groups (n = 3/group). Prior to surgery, the animals were allowed to adapt to the environment for at least one week. In this experiment, rats were anesthetized with 1 % pentobarbital sodium (50 mg/kg, i.p.), the tMCAO model was prepared by Zea Longa’s method(31). Throughout surgery, body temperature was maintained at 36.5 ± 0.5°C using a heating pad. Sham-operated rats underwent identical surgical procedures except for artery occlusion.

### 2.12. Humane endpoints, animal welfare, and monitoring

The planned experimental endpoint was 24 hours after reperfusion. To prevent severe suffering, predefined, objective humane endpoints were used. The specific criteria for immediate euthanasia were: (1) body weight loss exceeding 20% of the pre-surgical baseline; (2) complete inability to access food or water for 12 consecutive hours; (3) severe and persistent neurological deficits, specifically the loss of righting reflex for more than 30 seconds. Animal health and behavior were monitored at least every 4 hours during the immediate postoperative recovery period and at least twice daily thereafter. No animal met these humane endpoint criteria prior to the planned endpoint, and no unanticipated deaths occurred. All six animals (n=6) survived until the scheduled endpoint and were euthanized. To minimize pain and distress, the following measures were taken: precise intraoperative temperature control (maintained at 36.5 ± 0.5°C), provision of post-operative thermal support, ad libitum access to standard chow and water, and the provision of softened food and water on the cage floor post-surgery to facilitate intake. All personnel involved in animal procedures and monitoring received specific training on recognizing signs of rodent pain, distress, and the application of humane endpoint criteria.

### 2.13. TTC staining

After reaching the planned endpoint of 2 h of tMCAO and 24 h of reperfusion, the rats were deeply anesthetized with sodium pentobarbital (1%, 150 mg/kg, i.p.) and euthanized by decapitation while under deep anesthesia. The brain tissue was collected and immediately cut into five consecutive coronal sections (2 mm in thickness) with a slicing mold. The sections were subjected to staining with 0.5% TTC, washed in PBS, and then fixed in 4% paraformaldehyde for 6 hours. Finally, the infarction volume, expressed as a percentage, was analyzed using ImageJ software. The calculation was performed according to the following formula: Infarct volume (%) = (Contralateral hemisphere area - Non-infarcted ipsilateral hemisphere area) / (2 × Contralateral hemisphere area) × 100%.

### 2.14. Evaluation of OS activity in animal models

Extract brain tissues from rats in the sham surgery group and MCAO group on ice, weigh the extracted tissues, and add 0.9% physiological saline in a ratio of 1:9 weight to volume. At 4 ℃, centrifuge the tissue homogenate for 10 minutes and extract the supernatant. Measure the MDA level and SOD activity using the MDA detection kit (A003, Nanjing Jiancheng Bioengineering Institute) and total superoxide dismutase (T-SOD) detection kit (A001, Nanjing Jiancheng Bioengineering Institute).

### 2.15. Western blotting

The brain tissue proteins of three rats in the sham operation group and three rats in the MCAO group were extracted, 40 micrograms of protein were separated in 15% SDS-PAGE gel, and transferred to the polyvinylidene fluoride membrane. Seal the membrane with 5% skim milk powder at room temperature for 2 hours, then incubate overnight at 4 ° C with the following primary antibodies: Clusterin (*Clu*) rabbit monoclonal antibody (1:1000, ab69644, abcam), Glutamate Ionotropic Receptor AMPA Type Subunit 2 (*Gria2*) rabbit polyclonal antibody (1:1500, A11316, ABclonal Technology, China), and β-Tubulin rabbit polyclonal antibody (1:1500, A12289, ABclonal Technology, China). Subsequently, the membrane was incubated with a secondary antibody bound to polyvinylidene fluoride membrane and horseradish peroxidase at room temperature for 2 hours. Visualize Western blot using ECL chemiluminescence assay kit (P0018S, Beijing, China) and perform protein quantification using Image J.

### 2.16. Statistical Analysis

All data processing and statistical analysis were conducted using R version 4.4.3 and GraphPad Prism 9.0. The data is presented as mean ± standard error. The differences in continuous variables between two groups were evaluated using Wilcoxon test or t-test, while the differences between multiple groups were evaluated using one-way analysis of variance (ANOVA). The correlation between variables is evaluated using Pearson correlation coefficient analysis. All p-values were calculated using the two tailed method, with statistical significance defined as p-value < 0.05.

## 3. Results

### 3.1 Single cell analysis and identification of OS related genes

The workflow diagram of this study is shown in Figure 1. This study is based on the screening criteria of ribosome, mitochondrial, and red blood cell ratios. After a strict quality control process, the single-cell dataset was subjected to quality control screening, and a total of 27856 high-quality cells that meet the criteria were retained for further in-depth analysis. We performed batch correction on all samples, and the results showed that the cells in all samples were uniform, indicating successful correction of batch effects (Figure 2A). Through standard processing procedures, all cells were divided into 12 different cell clusters (Figure 2B). Through the joint annotation of classic marker genes and CellMarker 2.0, 11 different cell types were identified: Astrocytes, Endothelial cells, Ependymal cells (Fibroblasts, Macrophages, Microglia cells, Mural cells, Neuron cells, Neutrophil cells, Oligodendrocytes and T cells (Figure 2C-E), UMAP visualized the distribution of marker genes across cell populations (Figure S1). Based on the GO analysis of the 11 different cell types mentioned above, the enriched pathways can provide us with a deeper understanding of these cell categories (Figure 2C). In addition, different proportions of these 11 cell types were observed between the sham surgery group and the MCAO group (Figure 2F). In the MCAO group, the proportion of endothelial cells decreased, while the proportion of macrophages and mural cells increased. There was no significant difference in the proportion of other types of cells. To investigate the OS activity at the single-cell level, we used five algorithms including AUCell, Ucell, singlescore, ssgsea, and AddModuleScore to calculate the activity of each cell. The results showed heterogeneity in OS activity among different cell types, with the highest OS activity in mural cells and lower activity in T cells, endothelial cells, and astrocytes (Figure 3A). The UMAP results further demonstrated that mural cells have high OS activity (Figure 3B). According to the median OS score, cells were divided into high OS activity group and low OS activity group (Figure 3C). The analysis of differentially expressed genes between the high OS group and the low OS group identified a total of 308 related genes that affect OS activity (Figure 3D, Table S3), and the results of the correlation analysis identified 85 genes significantly associated with OS (Figure 3E, Table S4). Finally, a total of 36 genes overlapped between correlation analysis and differential expression analysis (Figure 3F, Table S5). We conducted GO analysis on these 36 genes to elucidate the functional roles (Figure 3G, Table S6). In the category of biological processes, these genes are associated with the differentiation of glial cells and cellular responses to brain-derived neurotrophic factor stimulation pathways. Cellular components are closely related to growth cone. In the molecular functional category, they are closely related to the activity of long-chain fatty acid Coa ligase. In addition, we also analyzed the expression differences of these 36 genes at the overall RNA-sep level between the sham group and the MCAO group (Figure S2).

**Figure 1.**
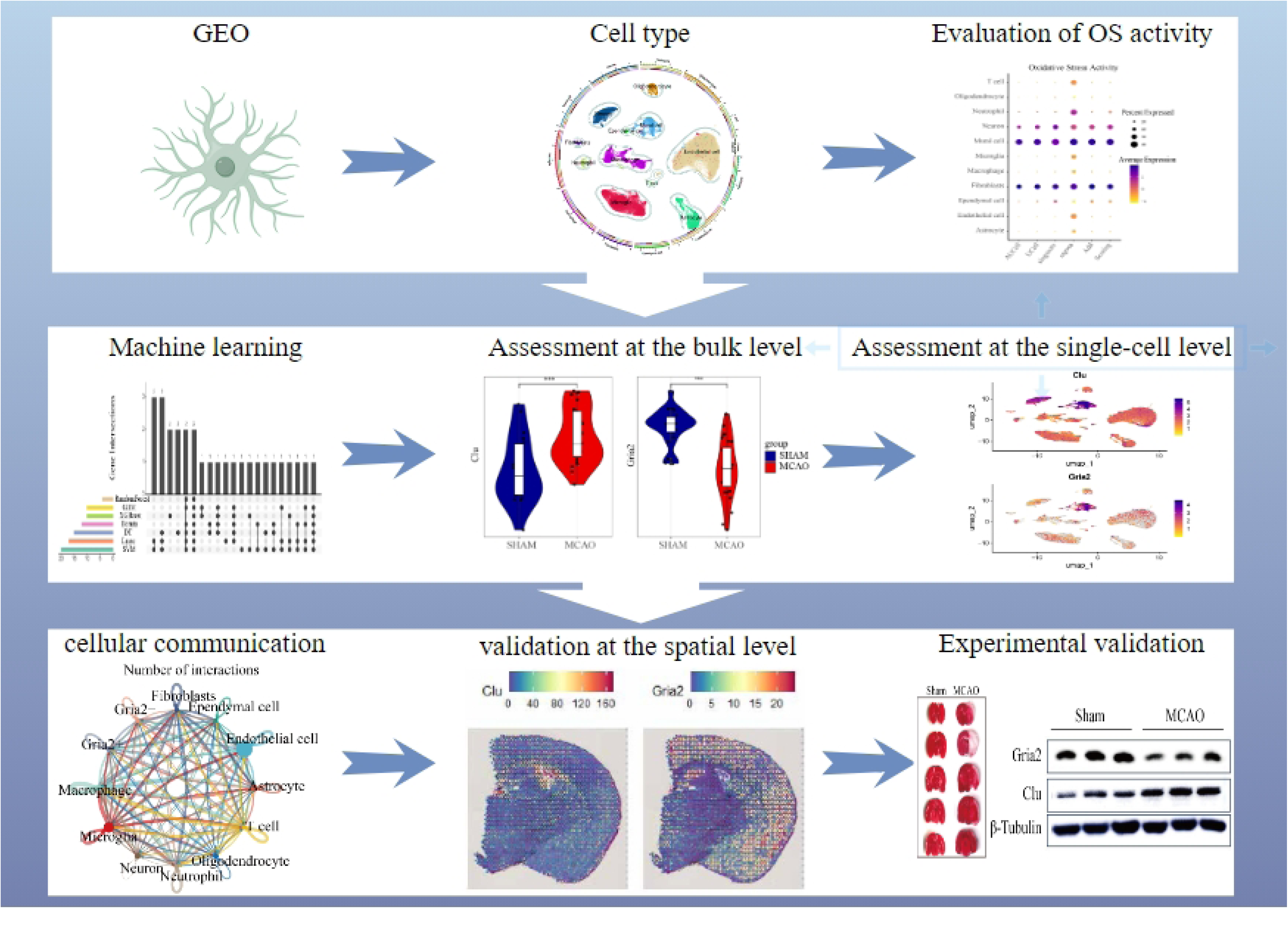
Flowchart depicting the study design.

**Figure 2.**
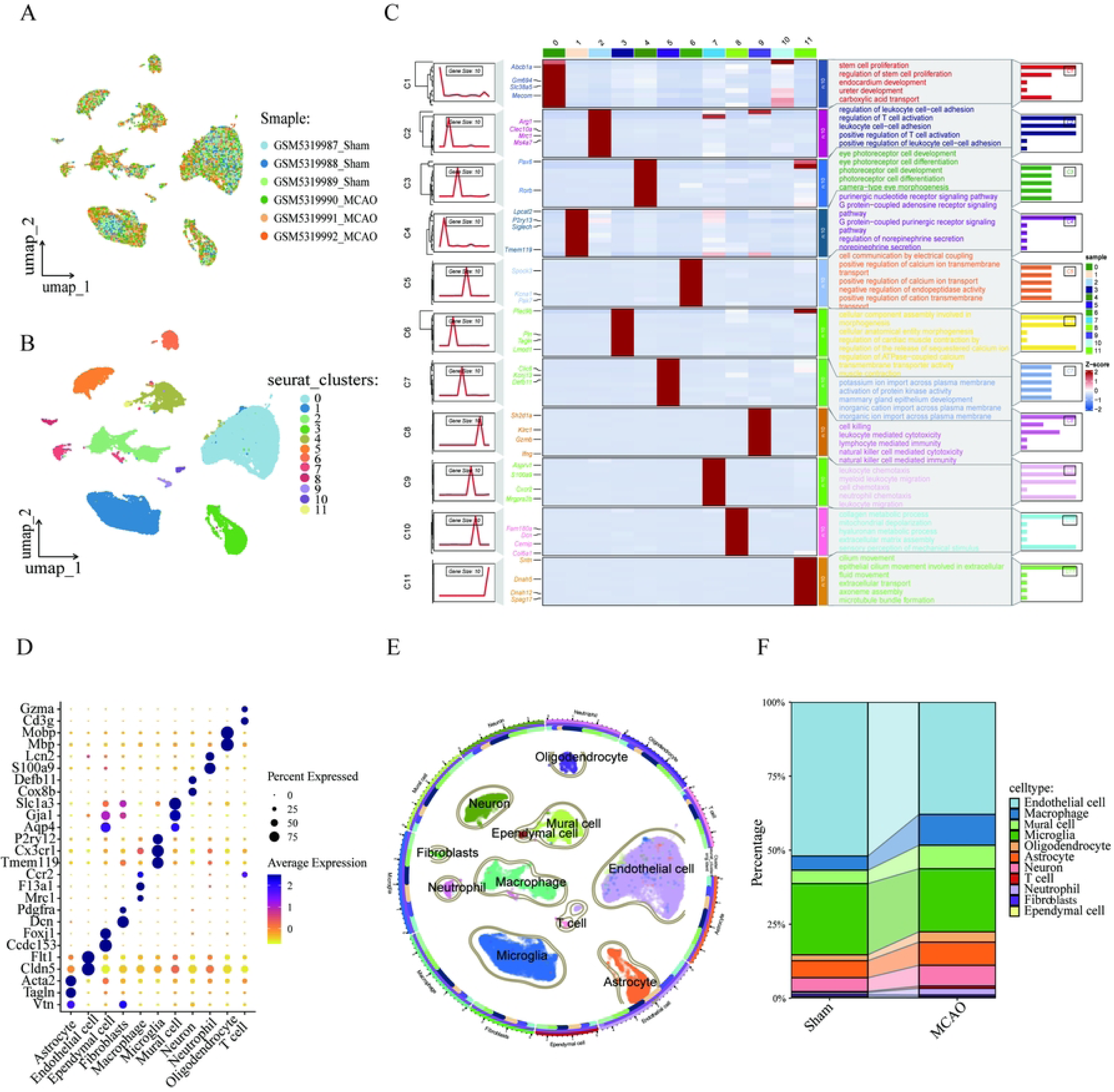
Screening of single-cell data. A. The PCA results revealed a relatively stable cell distribution across all analysed samples, with low sensitivity to batch effects. B. The results of the UMAP analysis indicated that all cells were meticulously classified into 12 clusters. C. The relationship between the marker genes of 12 clusters, along with the relevant pathways enriched by GO analysis. D–E. Based on classic marker genes, the data were manually annotated to 11 different cell types. F. Percentage of various cell types in the Sham and MACO groups.

**Figure 3.**
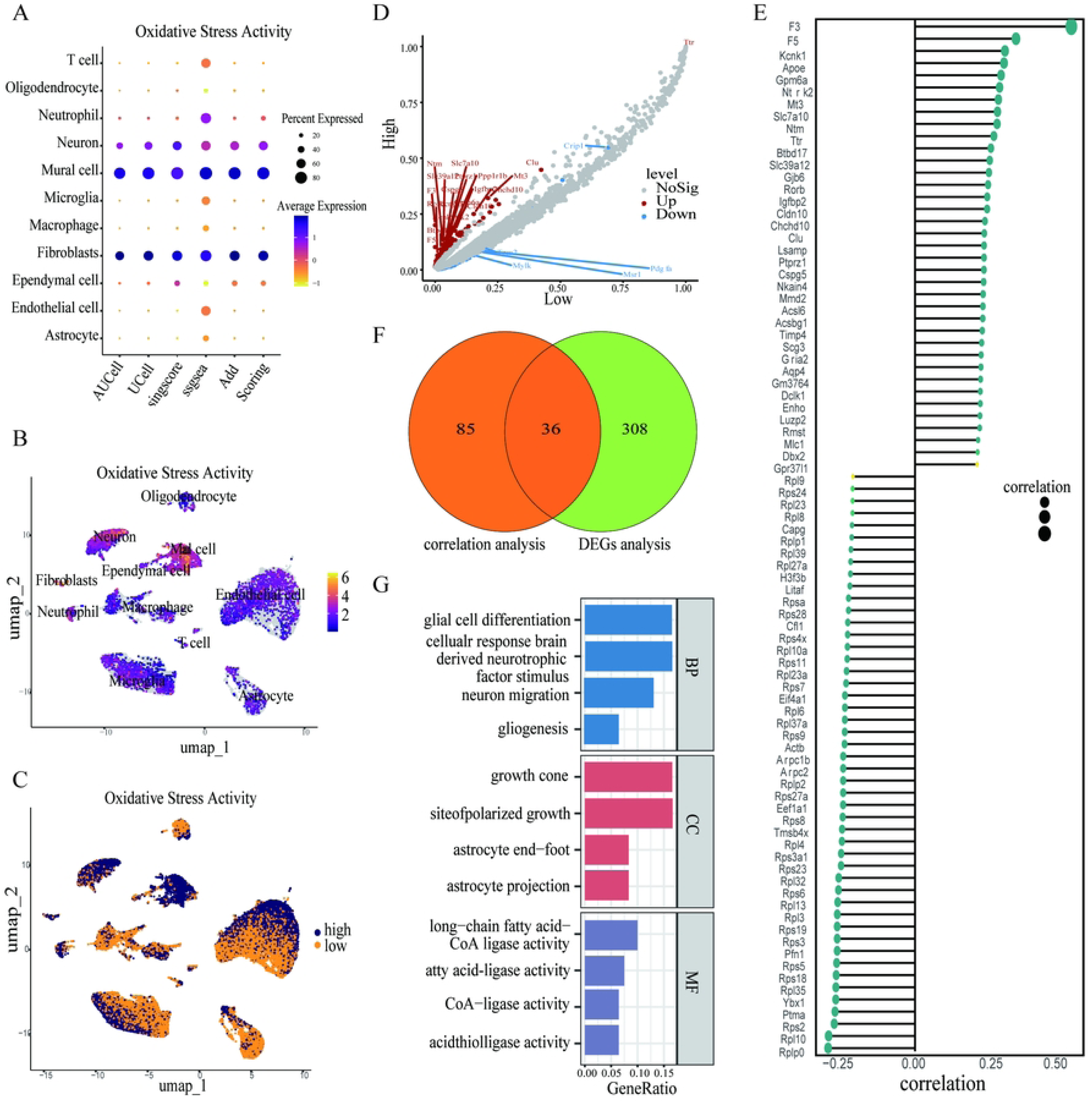
Analysis of single-cell OS activity in MCAO. The results of the AUCell, Ucell, singscore, ssgsea, and AddModuleScore algorithms showed that Mural cells had the highest aggregation activity. B. The results of the UMAP analysis indicated high OS activity in Mural cells. C. All cells were classified as high OS cells and low OS cells according to the median of the total OS activity. D. Results of DEGs analysis of OS. E. Correlation analysis was used to select genes that were significantly correlated with the OS scores. F. Results of Venn diagram of correlation analysis and DEGs analysis. G. The results of GO analysis of the OS genes.

### 3.2 Machine learning algorithms and bulk level validation indicate significant disease-specific expression of *Clu* and *Gria2*

We used seven machine learning algorithms to identify the best feature genes among 36 genes. After using the LASSO regression algorithm, we obtained 18 key genes (Figure 4A, Table S7); Through the random forest algorithm, we obtained 4 key genes (Figure 4B, Table S7), and after 5-fold cross validation using the SVM-RFE algorithm, we selected 20 key genes (Figure 4C, Table S7). The GBM algorithm identified 10 key genes (Figure 4D, Table S7). Fifteen key genes were obtained through the decision tree algorithm (Figure 4E, Table S7). Twelve key genes were obtained using the Boruta algorithm (Figure 4F-G, Table S7). The XGBoost algorithm ultimately identified 10 key genes (Figure S3, Table S7). Finally, after merging the key genes obtained from seven machine learning algorithms, we ultimately obtained two optimal feature genes: *Clu* and *Gria2*. (Figure 4H, Table S7). To verify the accuracy of the analysis, we evaluated these two optimal feature genes at the bulk RNA-seq level. In the GSE233811 dataset, compared to the Sham group, the MCAO group had higher levels of *Clu* expression and lower levels of *Gria2* (Figure 5A). The ROC curve shows that the AUC of *Clu* is 0.79 and the AUC of *Gria2* is 0.91 (Figure 5B). The correlation analysis between two genes showed a negative correlation (r=-0.55) between the expression level of *Clu* and *Gria2*. Afterwards, we further validated using the GSE137482 dataset, and found that *Clu* was highly expressed in the MCAO group, while *Gria2* was lowly expressed in the MCAO group (Figure 5D). The ROC curve shows that the AUC of *Clu* is 0.85 and the AUC of *Gria2* is 0.72. Correlation analysis shows that the two genes are still negatively correlated (r=-0.24), which confirms the robustness of the analysis.

**Figure 4.**
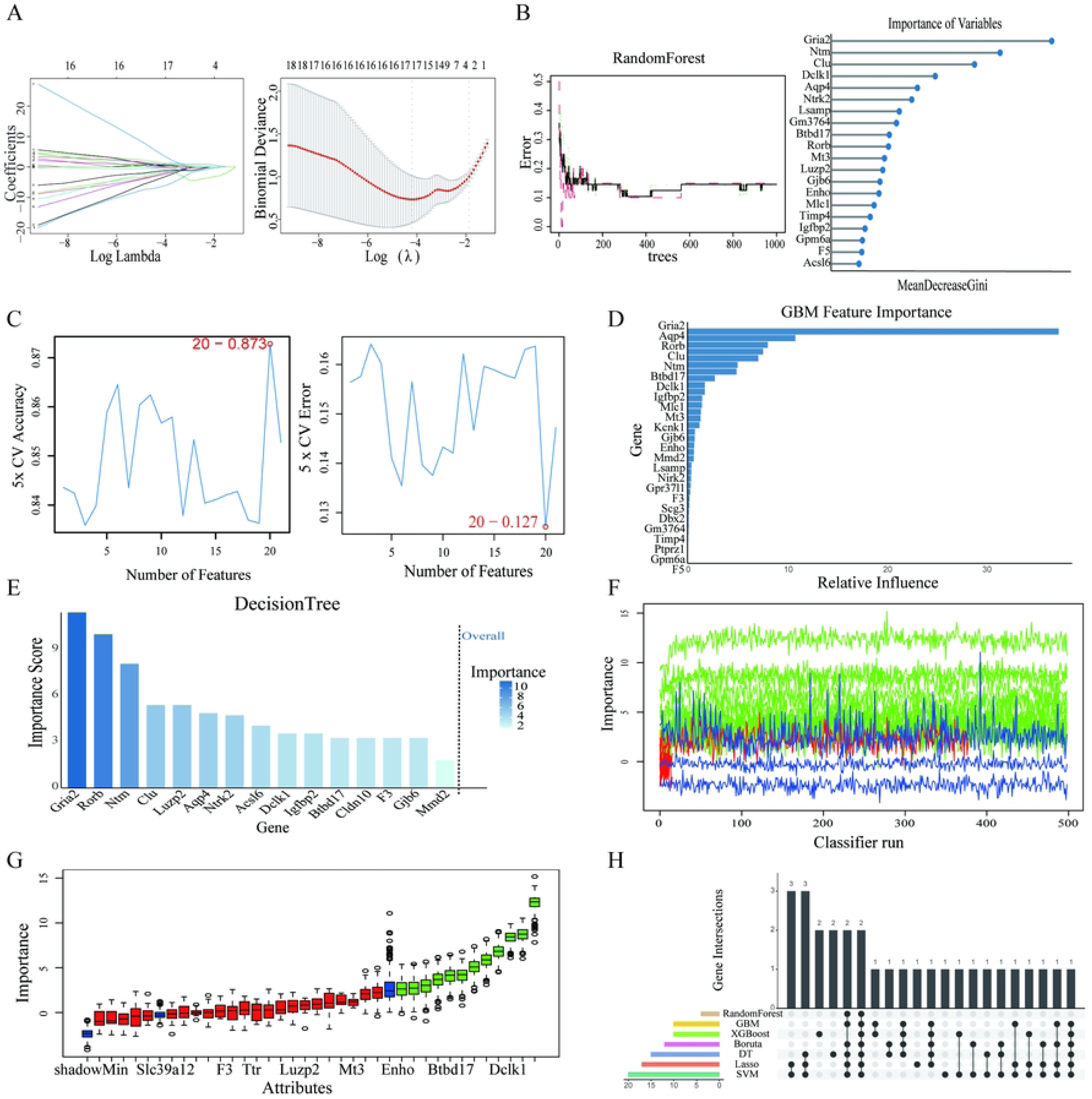
Seven machine learning algorithms integrated to identify optimal feature genes. A. Results of LASSO algorithm. B. Random forest (RF) for the relationship between the number of trees and error rate. C. Estimation of five fold cross-validation error using SVM-RFE. D–E. Estimation of fivefold cross-validation error using SVM-RFE. F. Influence of number of Decision Trees (DT) on error rate. G–H. Importance of features according to the Boruta algorithm. I. Venn diagram showing the seven optimal feature genes shared by LASSO, random forest, DT, GBM, XGBoost, Boruta, and SVM-REF algorithms.

**Figure 5.**
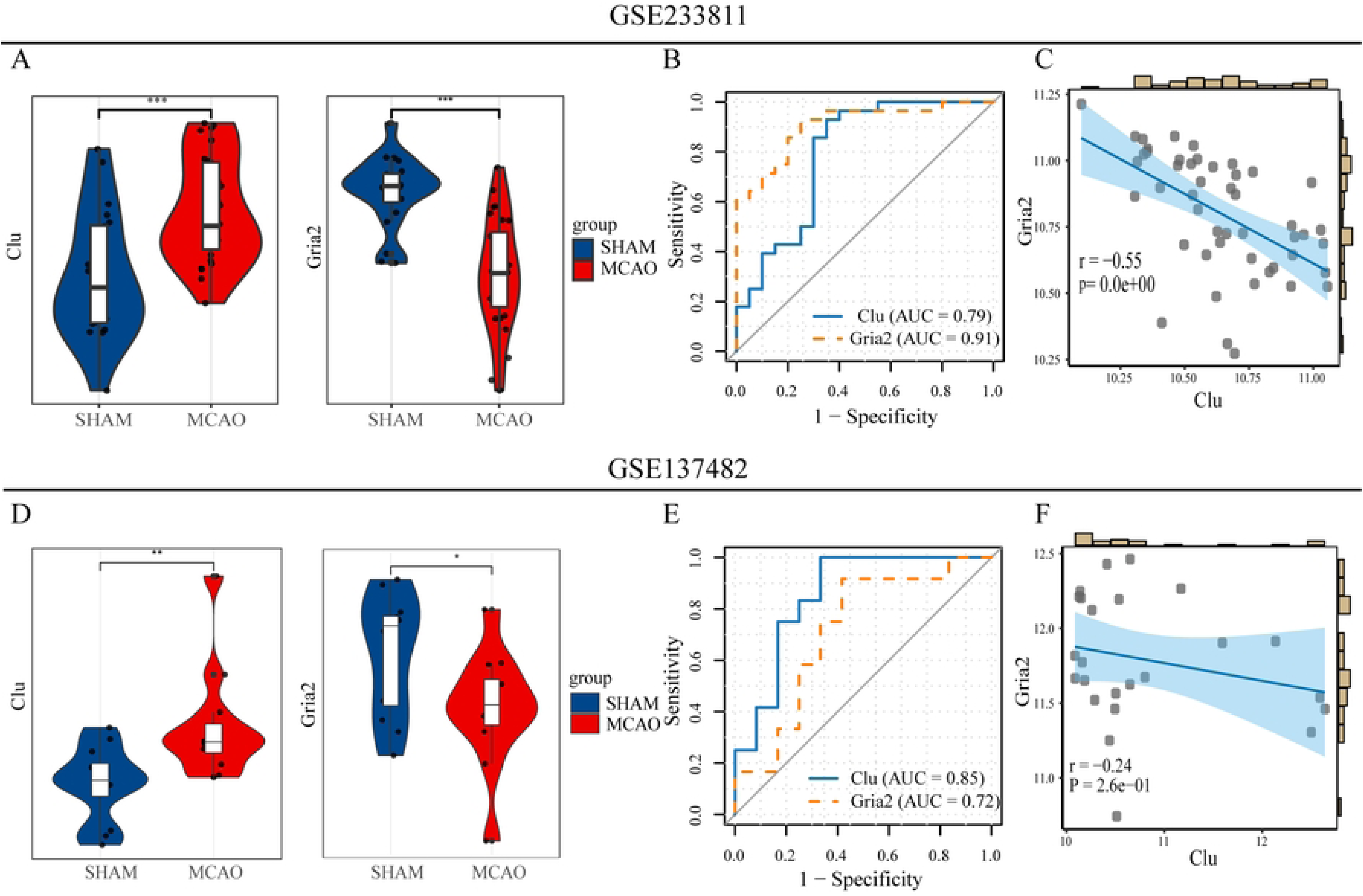
Validation of optimal feature genes at the bulk level. Expression levels of *Clu* and *Gria2* show significant differences between Sham and MCAO groups in GSE233811. B. The area under the curve indicates the diagnostic value of *Clu* and *Gria2*. C. The results of the Pearson correlation coefficient analysis indicated a notable correlation between the expression of *Clu* and *Gria2*. D. Expression levels of *Clu* and *Gria2* show significant differences between Sham and MCAO groups in GSE137482. E. The area under the curve indicates the diagnostic value of *Clu* and *Gria2*. F. The results of Pearson correlation coefficient analysis indicated a notable correlation between the expression of *Clu* and *Gria2*. Data are presented as mean ± SD, *\*p* < 0.05, *\*\* p* < 0.01 and *\*\*\* p* < 0.001 by Student’s two-tailed unpaired t-test.

### 3.3 *Clu* and *Gria2* are highly expressed in mural cells

We explored the specific cell types with optimal expression of characteristic genes at the single-cell level, and the results showed that *Clu* and *Gria2* were highly expressed in mural cells, with the lowest expression in T cells (Figure 6A). UMAP visualization shows that two genes are mainly located in mural cells (Figure 6B). This is consistent with our previous discovery of the highest OS activity in mural cells.

**Figure 6.**
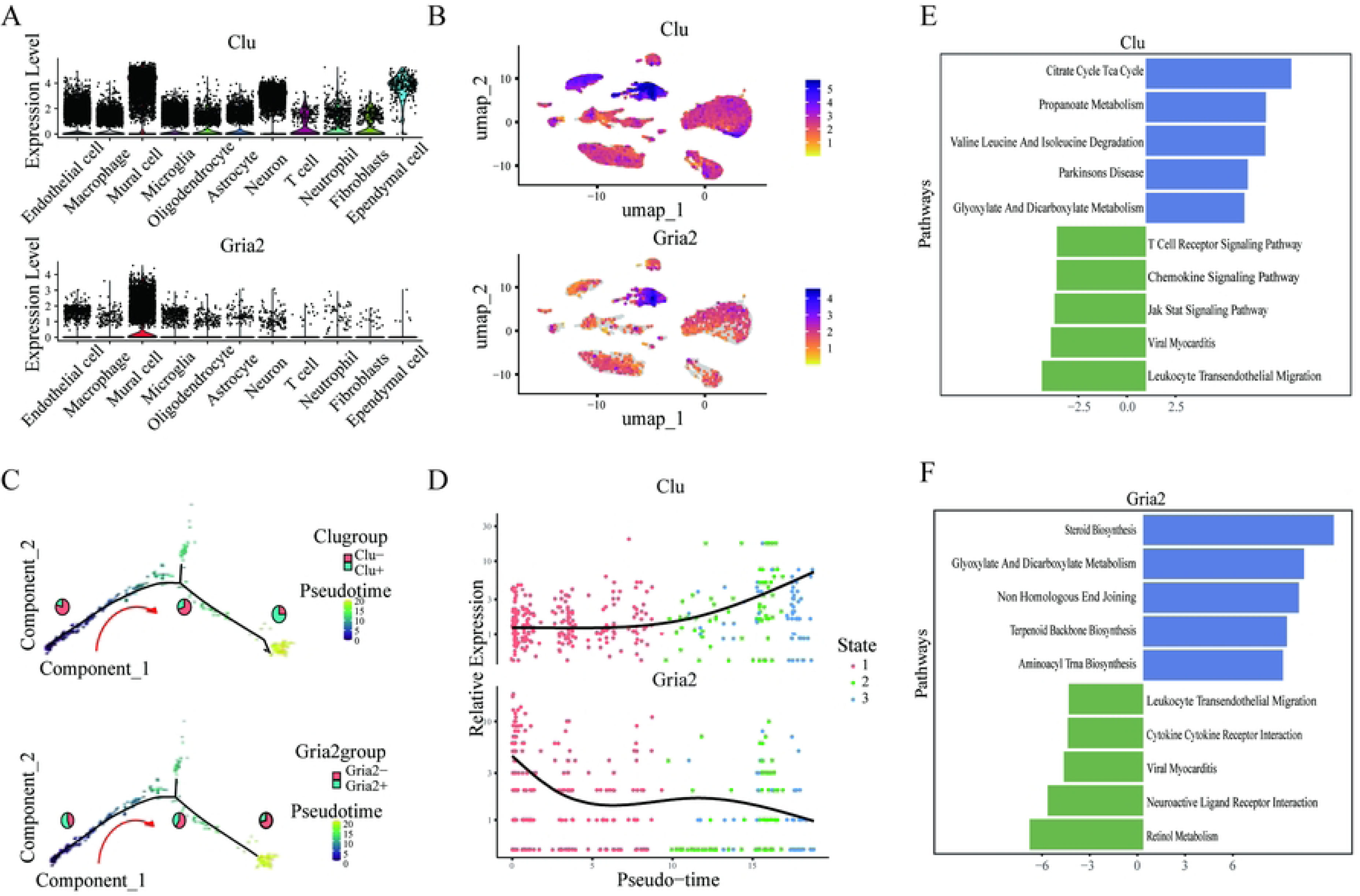
Validation of optimal feature genes at the single-cell level. A. The results of violin plot for *Clu* and *Gria2*. B. The results of UMAP indicated that *Clu* and *Gria2* were predominantly expressed in Mural cells. C. During the pseudotemporal differentiation trajectory, the relative abundances of *Clu*+ and *Gria2*+ Mural cells exhibited synchronized dynamic changes. D. The relative expression of *Clu* and *Gria2* in the pseudotemporal analysis. E. Results of GSVA Analysis of *Clu*-Associated Pathways. F. Results of GSVA Analysis of *Gria2*-Associated Pathways.

### 3.4 Trajectory analysis

We divide mural cells into *Clu*+mural cells and *Clu*-mural cells, as well as *Gria2*+mural cells and *Gria2*-mural cells. We explored transcriptional heterogeneity of mural cells through Monocle2 trajectory analysis. During the quasitemporal process, the proportion of *Clu*+mural cells increases, while the proportion of *Gria2*+mural cells decreases synchronously (Figure 6C). Pseudo time analysis shows the relative expression patterns of *Clu* and *Gria2* (Figure 6D).

### 3.5 GSVA results of *Clu* and *Gria2*

In order to explore the potential regulatory mechanisms of the optimal feature genes, we conducted GSVA analysis. The results showed that *Clu* was significantly enriched in citrate cycle Tca cycle, propionate metabolism, valine, leucine and isoleucine degradation, Parkinson’s disease, and oxalate and dicarboxylate metabolism, while there was relatively no enrichment in T cell receptor signaling pathway, chemokine signaling pathway, JAK-STAT signaling pathway, viral myocarditis and leukocyte trans-endothelial migration (Figure 6E, Table S8). On the other hand, *Gria2* is significantly enriched in steroid biosynthetic, glycoxylate and dicarboxylate metabolism, non-homologous end joining, terpenoid backbone biosynthetic and aminocyclic trna biosynthetic pathways, while it was relatively no enrichment in leukocyte transendothelial migration, cytokine cytokine receptor interaction, viral myocarditis, neuroactive ligand receptor interaction and retinol metabolism (Figure 6F, Table S9).

### 3.6 Results of cell interactions

The results of intercellular communication showed the quantity and intensity of interactions between *Clu*+, *Clu*-, *Gria2*+, and *Gria2*-mural cells and other cell types (Figure 7A-B). Compared with *Clu*-mural cells, *Clu*+ mural cells exhibited higher input and output intensity, and similarly, *Gria2*+ mural cells also had higher input and output intensity than *Gria2*-mural cells (Figure 7C). The ligand receptor interaction analysis demonstrated the interactions between different cell types and *Clu*+, *Clu*-, *Gria2*+, and *Gria2*-mural cells (Figure 7D-E). The results showed that *Clu*+ mural cells mainly interact with other cells through Ptn-Ncl and Mdk-Ncl receptor ligands. *Gria2*+ mural cells mainly interact with other cells through Vegfa-Vegfr1, Ptn-Ncl, and Mdk-Ncl receptor ligands.

**Figure 7.**
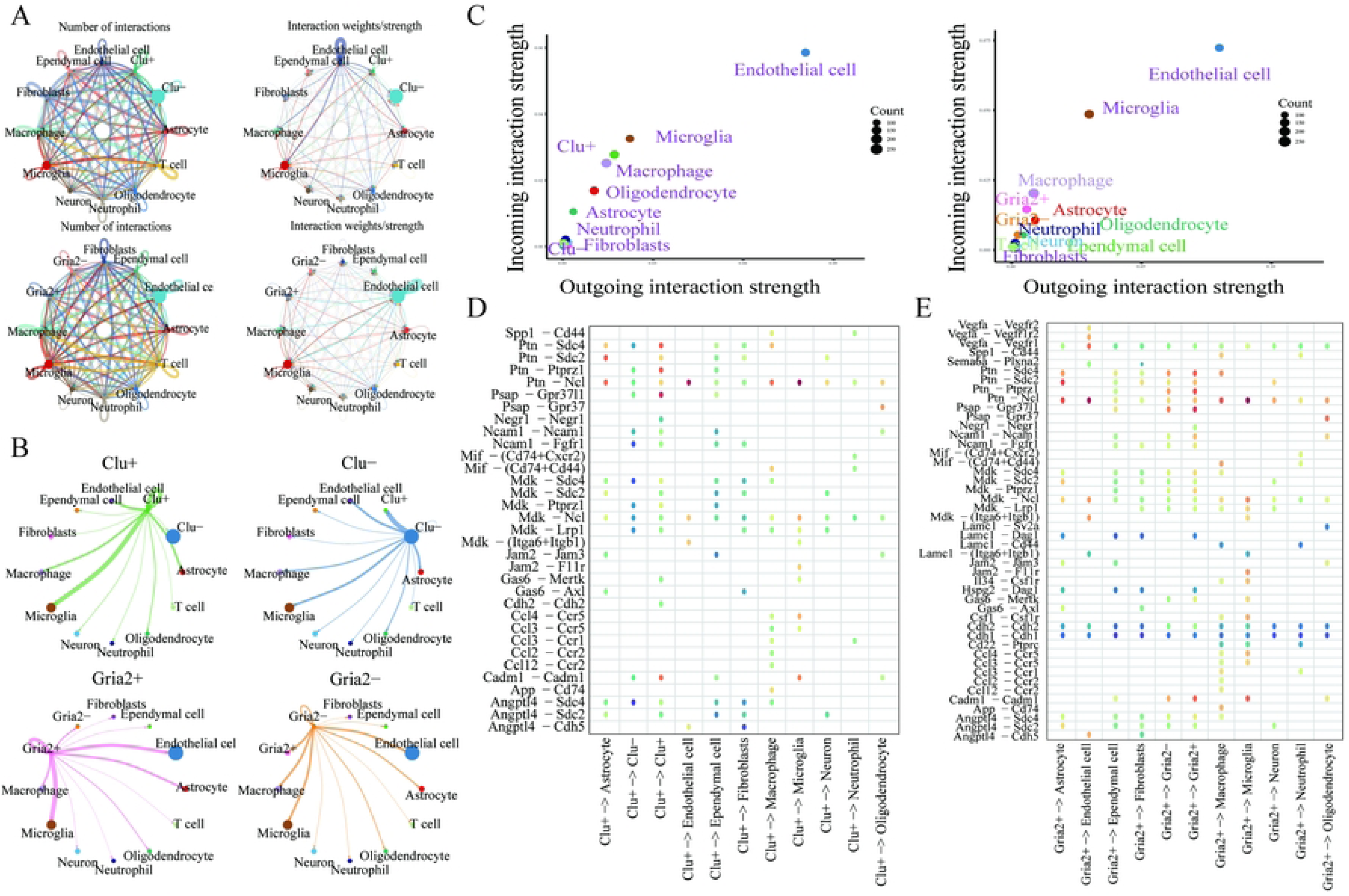
Cell interaction analysis. A-B. The results reveal the cellular interaction weights and the number of interactions between *Clu*+ and *Clu*− Mural cells, *Gria2*+ and *Gria2*− Mural cells, as well as other cell types within the microenvironment of MCAO. C. The relationship between differential outgoing interactions and incoming interaction strength is examined in *Clu*+ Mural cells, *Clu*− Mural cells, *Gria2*+ Mural cells, and *Gria2*− Mural cells. D-E. Ligand-receptor interactions between different cell types and *Clu*+ Mural cells, as well as *Gria2*+ Mural cells.

### 3.7 High expression of *Clu* and *Gria2* in mural cells at the ST level

On Sham slices, a total of 13.9 million UMIs and 32285 genes were detected in 2417 spots, with approximately 5800 UMIs and 1400 unique genes detected in one spot. On MCAO slices, a total of 9.6 million UMIs and 33538 genes were detected in 2587 spots, and approximately 3800 UMIs and 1300 unique genes were detected in one spot. We analyzed and integrated ST data and scRNA-seq data. The Sham group identified 5 cell types: Mural cells, Oligodendrocytes, Neuron cells, Fibroblasts, and Ependymal cells. And The MCAO group obtained a total of 10 cell types: Ependymal cells (Oligodendrocytes, Neuron cells, Fibroblasts (Macrophages, Neutrophil cells, Microglia cells, Mural cells, Astrocytes And Endothelial cells (Figure 8A-D). We explored the spatial expression distribution of *Clu* and *Gria2*, and the results were consistent with the single-cell level. Both genes were highly expressed in parietal cells (Figure 8E-F).

**Figure 8.**
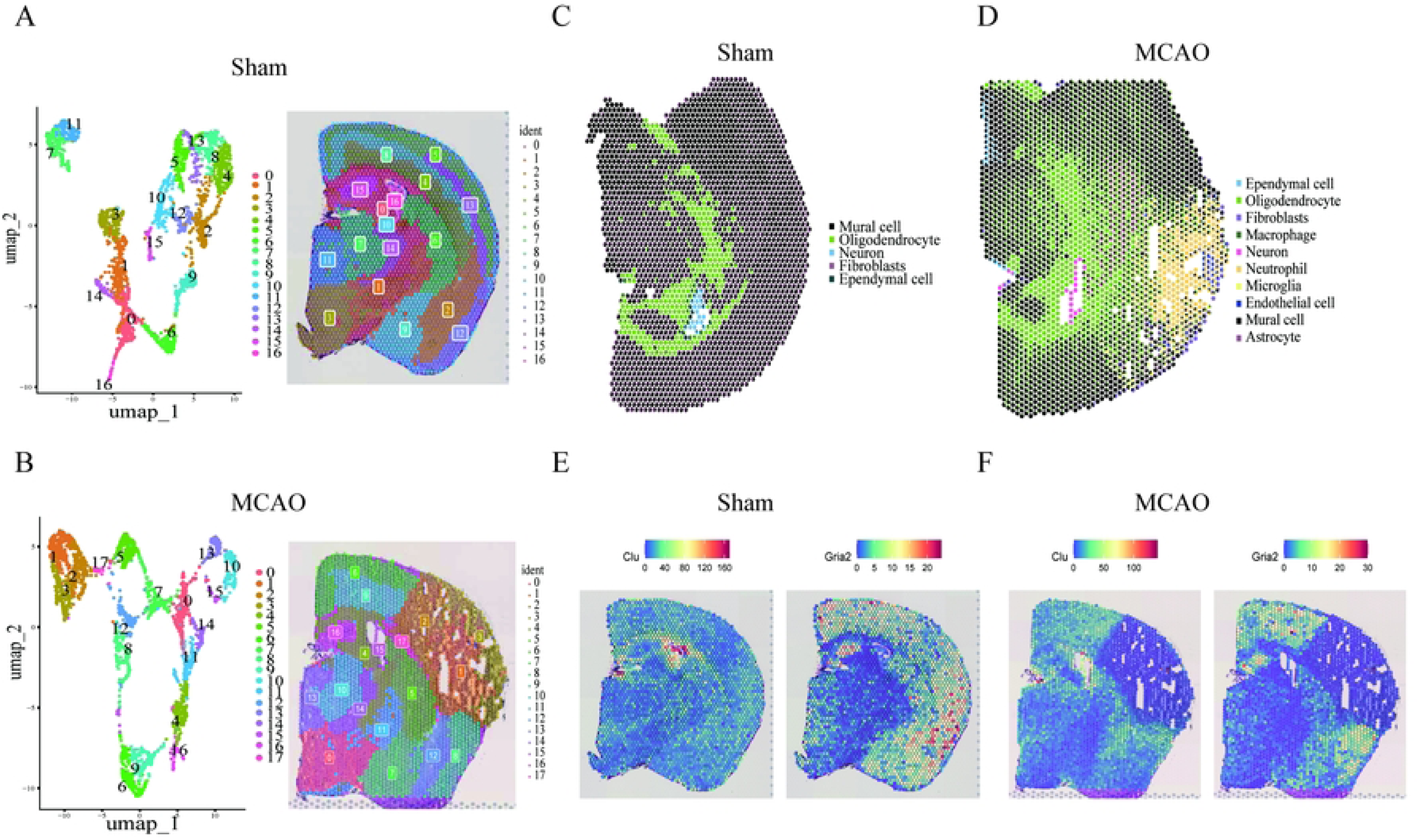
optimal feature genes at the spatial level. A. UMAP plot of Sham ST spots(left) and mapped with unbiased clustering of ST spots in Sham sections (right). B. UMAP plot of MACO ST spots(left) and mapped with unbiased clustering of ST spots in MCAO sections (right). C-D. Spatial plot of deconvoluted cell type proportions per spot in Sham and MCAO sections. E. The spatial location and gene changes of *Clu* and *Gria2* were visualized in Sham sections. F. The spatial location and gene changes of *Clu* and *Gria2* were visualized in MCAO sections.

### 3.8 Experimental verification of *Clu* and *Gria2*

Firstly, TTC staining was performed on rat brain slices, with undamaged brain tissue areas stained red and damaged tissue areas stained white. Significant infarcted areas were observed in the MCAO group (Figure 9A). Afterwards, we detected the expression levels of malondialdehyde (MDA) and total superoxide dismutase (T-SOD) in two groups of brain tissues. In the MCAO group, the expression level of MDA significantly increased while the expression level of T-SOD significantly decreased, indicating higher oxidative stress activation in the MCAO group (Figure 9B). The results of western blotting showed that, compared with the Sham group, the protein expression of *Clu* increased in the MCAO group, while the expression of *Gria2* decreased (Figure 9C-D).

**Figure 9.**
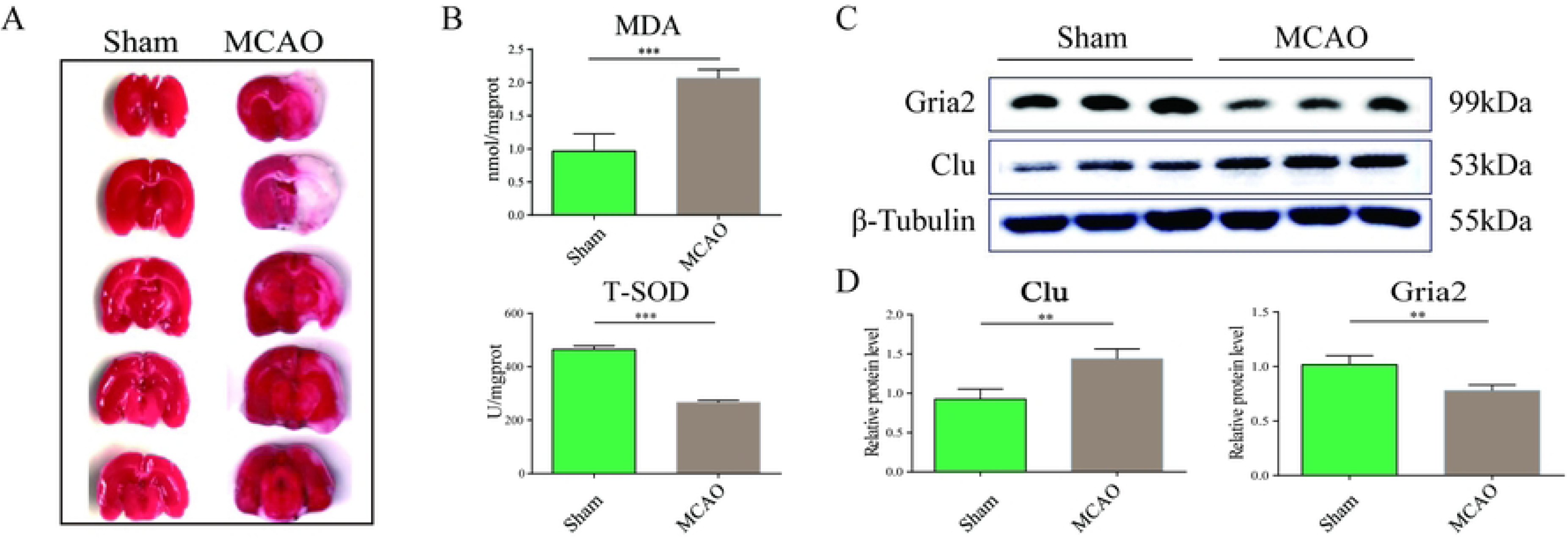
Results of experimental verification of the rat model. A. The effects of cerebral ischemia-reperfusion injury in rats by dyeing results of TTC. B. There was a significant increase in the expression of MDA in HF tissues, accompanied by a decreased expression of T-SOD. n = 3/group. C-D. The results of western blotting revealed that compared to the sham group, the protein expression of *Clu* increased and the protein expression of *Gria2* decreased in the MCAO group.

## 4. Discussion

Stroke is an acute cerebrovascular disease that occurs rapidly and progresses rapidly, with over 13.7 million new strokes occurring worldwide each year. IS is the most common type of stroke, accounting for about 80% of the total number of strokes (32). After IS, neuronal loss caused by ischemia and infarction is the most direct cause of neuronal damage. Excitatory toxicity, OS and mitochondrial dysfunction caused by ischemia and hypoxia can lead to a large number of neuronal apoptosis (33). Currently, an increasing number of studies have shown that OS is one of the key factors contributing to secondary brain injury after IS (34, 35). In this study, we found that the highest OS activity was observed in mural cells through a combination of scRNA-seq and machine learning. Additionally, we identified *Clu* and *Gria2* as key genes closely associated with OS in IS, primarily expressed in mural cells. To ensure the robustness of our findings, we conducted ST analysis and animal experiments to verify the expression levels of *Clu* and *Gria2* in IS. These results highlight the potential of these genes as therapeutic targets for future interventions aimed at modulating oxidative stress activity and mitigating ischemic stroke damage.

The pathophysiology of IS is a complex process, where free radicals and ROS increase in brain tissue, while antioxidant enzymes become inactive and protective antioxidant activity levels decrease, resulting in the inability of natural defense mechanisms to effectively protect neurons (36). OS, along with its associated inflammatory responses, autophagy, and apoptosis, plays a crucial role in neurodegenerative damage. Under ischemic conditions, the affected region experiences reduced blood flow, which leads to decreased oxygen partial pressure and elevated carbon dioxide partial pressure. These changes result in tissue acidosis, bioenergetic insufficiency, and the initiation of OS. These changes further damage cerebral microvessels and the blood-brain barrier, and activate neuroglial cells (37). Our research found that mural cells exhibit significantly higher OS activity than other types of cells. Mural cells are mainly located in the vascular wall, including pericytes and vascular smooth muscle cells. These cells cover the outer side of endothelial cells and are embedded in the basement membrane of blood vessels, playing a role in stabilizing vascular structure and regulating vascular function (38). As a critical component of the blood-brain barrier, the dysfunction of mural cells directly drives the deterioration of ischemic injury. During cerebral ischemia, excessive production of ROS and reactive nitrogen species occurs. Apoptosis or contraction of mural cells can lead to the disruption of the blood-brain barrier. Following ischemia, the accumulation of ROS in mural cells triggers abnormal contraction, resulting in the phenomenon of “no reflow” in capillaries and impeding the recovery of blood flow (39, 40). Moreover, after ischemia, mural cells release pro-inflammatory cytokines, which recruit neutrophils and monocytes to infiltrate the affected area. These cells further amplify ROS production through NADPH oxidase, creating a positive feedback loop of “OS-inflammatory infiltration”. This loop activates the NF-κB pathway, intensifying the inflammatory cascade reaction. Ultimately, this leads to microcirculatory collapse and secondary brain tissue damage (41–43). Therefore, antioxidant therapy targeting mural cells may provide a new direction for reversing ischemic vascular injury.

We identified key genes with strong diagnostic potential: *Clu* and *Gria2*, by integrating single-cell sequencing data and applying machine learning algorithms. Afterwards, the diagnostic value of *Clu* and *Gria2* was further supported by ROC curve analysis through bulk RNA-seq dataset validation. Both genes showed high AUC values, highlighting their predictive accuracy for IS diagnosis. *Clu* is a multifunctional glycoprotein widely present in various cellular components, widely involved in various biological processes such as cell protection, lipid metabolism, cell adhesion and migration, immune regulation, neuroprotection, tumor occurrence and metastasis. It plays an important role in various diseases such as Alzheimer’s disease, Parkinson’s disease, fibrosis, metabolic diseases, cardiovascular and cerebrovascular diseases, and tumors (44, 45). Research has demonstrated that elevated levels of oxidative stress are associated with increased *Clu* expression in asthma patients (46). Similarly, *Clu* expression is significantly upregulated during renal and cardiac ischemia-reperfusion (47, 48). Experimental evidence indicates that in cerebral ischemia models, increased *Clu* expression is significantly correlated with reduced infarct volume and decreased levels of oxidative stress markers, such as malondialdehyde and 8-hydroxy-2’-deoxyguanosine. Furthermore, exogenous supplementation with *Clu* has been shown to effectively reduce intracellular ROS accumulation and enhance cell survival rates (49, 50). These findings are consistent with our observations. After ischemic injury, the expression of *Clu* is significantly upregulated, which may be a cellular protective mechanism. As a molecular chaperone, *Clu* can bind to misfolded proteins, preventing their aggregation and formation of toxic amyloid fibers, thereby reducing cellular damage caused by OS. In addition, *Clu* enhances the antioxidant and survival abilities of cells by regulating the PI3K Akt and GSK-3 β signaling pathways (49, 51). Based on these findings, *Clu* may emerge as a promising therapeutic target. However, additional research is required to uncover the precise mechanisms by which *Clu* influences OS in IS.

*Gria2*, also known as *GluR2*, is a key subunit of the AMPA-type glutamate receptor and belongs to the family of ionotropic glutamate receptors. Downregulation of *Gria2* has been observed in various types of neurologic injuries and chronic neurodegenerative diseases, including ischemic-hypoxic brain injury, epilepsy, traumatic brain injury, Alzheimer’s disease, and amyotrophic lateral sclerosis (52). The expression level of *Gria2* determines the permeability of AMPA receptors to calcium ions. Research has shown that downregulation of *Gria2* expression during ischemia-reperfusion can lead to increased permeability of AMPA receptors to calcium ions, resulting in calcium overload and ultimately leading to cell death (53). Research has shown that electroacupuncture preconditioning can activate the cannabinoid CB_1_ receptor (CB_1_R), which in turn upregulates the expression of *Gria2*. This mechanism effectively reduces calcium ion influx, inhibits apoptosis, and enhances neurological function (54). Currently, direct studies investigating the relationship between *Gria2* and OS following cerebral ischemia are lacking. However, existing research has shown that modulating the expression or function of *Gria2* can improve neurological function and alleviate OS (55). Therefore, more research is needed in the future to help us better understand the specific mechanisms of *Clu* and *Gria2* in OS during IS.

This study has the following limitations. Firstly, model construction relies on public datasets, which may introduce dataset variability and patient selection related biases, although we try to minimize these biases through various machine learning algorithms and databases. In the future, the results can be further validated through more diverse cohort studies and in vitro or in vivo experiments. Secondly, although the correlation between OS activation and the expression of *Clu* and *Gria2* has been confirmed, its exact molecular mechanism in IS is still unclear. Future research needs to focus on revealing the signaling pathways and cellular processes regulated by *Clu* and *Gria2* under OS, and exploring the roles of their post transcriptional modifications (such as phosphorylation and ubiquitination) and post-translational modifications (such as glycosylation) in the pathogenesis of cerebral ischemia. In addition, although animal models have validated the correlation between these genes and IS, their diagnostic and therapeutic potential in humans still needs to be further validated through clinical studies. Future research needs to adopt a broader and more comprehensive approach to elucidate its mechanisms. Despite the limitations mentioned above, this study provides important insights into the molecular basis of IS, identifies potential diagnostic biomarkers and therapeutic targets, lays the foundation for future research, and provides beneficial directions for the development of diagnostic and therapeutic strategies for cerebral ischemia.

## 5. Conclusion

In this study, we redefined the gene set associated with OS in IS through scRNA-seq analysis. By integrating these findings with bulk RNA-seq data and employing machine learning algorithms, we identified two key genes: *Clu* and *Gria2*. The expression of *Clu* and *Gria2* can regulate OS activity, mainly in parietal cells. *Clu* and *Gria2* may participate in the progression of IS by altering the OS activity of parietal cells. These findings provide important evidence for improving the understanding of the role of OS in IS and provide reference for personalized treatment of the disease.

## Abbreviations

OS: Oxidative stress
IS: Ischemic stroke
ROS: reactive oxygen species
ScRNA-seq: Single-cell RNA sequencing
ST: Spatial transcriptomics
LASSO: Least Absolute Shrinkage and Selection Operator
GEO: Gene Expression Omnibus
MCAO: middle-cerebral artery occlusion
PCA: Principal component analysis
DEGs: Differential expressed genes
GO: Gene Ontology
ROC: Receiver operating characteristic
AUC: Area under the curve *Clu* Clusterin
MDA: Malondialdehyde
T-SOD: Total superoxide dismutase

## Clinical trial number

Not applicable.

## Data availability

The datasets utilised in this study, GSE174574, GSE233811, GSE137482 and GSE233813, were procured from the GEO database. All data and codes necessary for the analyses are available upon request. Please contact the corresponding author for further information.

## Acknowledgements

We are deeply grateful to all study participants and their families for their invaluable contributions.

## CRediT authorship contribution statement

Jiaxing Zhu: Writing - original draft, Project administration, Data curation, Conceptualization. Chunyan Zhou: Methodology, Investigation. Jing Yao: Data curation, Methodology. Jiali Xie: Writing - review & editing, Methodology, Formal analysis, Investigation, Funding acquisition.

## Funding

This work was supported by the grants from Jiangxi Provincial Administration of Traditional Chinese Medicine Science and Technology Plan (No. SZYYB20224260), Science and Technology Research Project of Jiangxi Provincial Department of Education (No. GJJ2209718), and Ganzhou City Guided Science and Technology Plan Project (No. GZ2024ZSF877).

## Declaration of competing interest

The authors declare that they have no known competing financial interests or personal relationships that could have appeared to influence the work reported in this paper.

## Supporting information

Fig. S1. UMAP visualized the distribution of marker genes across cell populations.

Fig. S2. the expression differences of these 36 genes at the overall RNA-sep level between the sham group and the MCAO group.

Fig. S3. The XGBoost algorithm.

**Table S1.** 2054 high confidence OS related genes were screened by setting the OS-related score threshold to ≥7.

**Table S2.** The typical marker genes.

**Table S3.** 308 related genes that affect OS activity between the high OS group and the low OS group.

**Table S4.** The results of the correlation analysis identified 85 genes significantly associated with OS.

**Table S5.** 36 genes overlapped between correlation analysis and differential expression analysis.

**Table S6.** GO analysis on these 36 genes to elucidate the functional roles.

**Table S7.** The results of the Machine learning algorithms.

**Table S8.** The results of the GSVA analysis of *Clu*.

**Table S9.** The results of the GSVA analysis of *Gria2*.

## Notes

### Competing Interest Statement

The authors have declared no competing interest.

## References

1. Daniele SG, Trummer G, Hossmann KA, Vrselja Z, Benk C, Gobeske KT, et al. Brain vulnerability and viability after ischaemia. Nat Rev Neurosci. 2021;22(9):553–72.

2. Chen H, Yoshioka H, Kim GS, Jung JE, Okami N, Sakata H, et al. Oxidative stress in ischemic brain damage: mechanisms of cell death and potential molecular targets for neuroprotection. Antioxidants & redox signaling. 2011;14(8):1505–17.

3. Li Z, Bi R, Sun S, Chen S, Chen J, Hu B, et al. The Role of Oxidative Stress in Acute Ischemic Stroke-Related Thrombosis. Oxidative medicine and cellular longevity. 2022;2022:8418820.

4. Rodrigo R, Fernández-Gajardo R, Gutiérrez R, Matamala JM, Carrasco R, Miranda-Merchak A, et al. Oxidative stress and pathophysiology of ischemic stroke: novel therapeutic opportunities. CNS & neurological disorders drug targets. 2013;12(5):698–714.

5. Jurcau A, Ardelean AI. Oxidative Stress in Ischemia/Reperfusion Injuries following Acute Ischemic Stroke. Biomedicines. 2022;10(3).

6. Li P, Stetler RA, Leak RK, Shi Y, Li Y, Yu W, et al. Oxidative stress and DNA damage after cerebral ischemia: Potential therapeutic targets to repair the genome and improve stroke recovery. Neuropharmacology. 2018;134(Pt B):208–17.

7. Ochocka N, Segit P, Walentynowicz KA, Wojnicki K, Cyranowski S, Swatler J, et al. Single-cell RNA sequencing reveals functional heterogeneity of glioma-associated brain macrophages. Nature communications. 2021;12(1):1151.

8. Jovic D, Liang X, Zeng H, Lin L, Xu F, Luo Y. Single-cell RNA sequencing technologies and applications: A brief overview. Clinical and translational medicine. 2022;12(3):e694.

9. Zhang L, Chen D, Song D, Liu X, Zhang Y, Xu X, et al. Clinical and translational values of spatial transcriptomics. Signal transduction and targeted therapy. 2022;7(1):111.

10. Guo W, Zhou B, Yang Z, Liu X, Huai Q, Guo L, et al. Integrating microarray-based spatial transcriptomics and single-cell RNA-sequencing reveals tissue architecture in esophageal squamous cell carcinoma. EBioMedicine. 2022;84:104281.

11. Greener JG, Kandathil SM, Moffat L, Jones DT. A guide to machine learning for biologists. Nature reviews Molecular cell biology. 2022;23(1):40–55.

12. Guan S, Xu Z, Yang T, Zhang Y, Zheng Y, Chen T, et al. Identifying potential targets for preventing cancer progression through the PLA2G1B recombinant protein using bioinformatics and machine learning methods. International journal of biological macromolecules. 2024;276(Pt 1):133918.

13. Sun X, Huang X, Sun X, Chen S, Zhang Z, Yu Y, et al. Oxidative Stress-Related lncRNAs Are Potential Biomarkers for Predicting Prognosis and Immune Responses in Patients With LUAD. Frontiers in genetics. 2022;13:909797.

14. Hao Y, Stuart T, Kowalski MH, Choudhary S, Hoffman P, Hartman A, et al. Dictionary learning for integrative, multimodal and scalable single-cell analysis. Nature biotechnology. 2024;42(2):293–304.

15. Aibar S, González-Blas CB, Moerman T, Huynh-Thu VA, Imrichova H, Hulselmans G, et al. SCENIC: single-cell regulatory network inference and clustering. Nature methods. 2017;14(11):1083–6.

16. Andreatta M, Carmona SJ. UCell: Robust and scalable single-cell gene signature scoring. Computational and structural biotechnology journal. 2021;19:3796–8.

17. Bhuva DD, Foroutan M, Xie Y, Lyu R, Cursons J, Davis MJ. Using singscore to predict mutation status in acute myeloid leukemia from transcriptomic signatures. F1000Research. 2019;8:776.

18. Jin Y, Wang Z, He D, Zhu Y, Chen X, Cao K. Identification of novel subtypes based on ssGSEA in immune-related prognostic signature for tongue squamous cell carcinoma. Cancer medicine. 2021;10(23):8693–707.

19. Mei Y, Li M, Wen J, Kong X, Li J. Single-cell characteristics and malignancy regulation of alpha-fetoprotein-producing gastric cancer. Cancer medicine. 2023;12(10):12018–33.

20. Fan C, Chen F, Chen Y, Huang L, Wang M, Liu Y, et al. irGSEA: the integration of single-cell rank-based gene set enrichment analysis. Briefings in bioinformatics. 2024;25(4).

21. Xu S, Hu E, Cai Y, Xie Z, Luo X, Zhan L, et al. Using clusterProfiler to characterize multiomics data. Nature protocols. 2024;19(11):3292–320.

22. Liang D, Wang L, Zhong P, Lin J, Chen L, Chen Q, et al. Perspective: Global Burden of Iodine Deficiency: Insights and Projections to 2050 Using XGBoost and SHAP. Advances in nutrition (Bethesda, Md). 2025;16(3):100384.

23. Hu J, Szymczak S. A review on longitudinal data analysis with random forest. Briefings in bioinformatics. 2023;24(2).

24. Dai W, Zheng P, Wu J, Chen S, Deng M, Tong X, et al. Integrated analysis of single-cell RNA-seq and chipset data unravels PANoptosis-related genes in sepsis. Frontiers in immunology. 2023;14:1247131.

25. Sanz H, Valim C, Vegas E, Oller JM, Reverter F. SVM-RFE: selection and visualization of the most relevant features through non-linear kernels. BMC bioinformatics. 2018;19(1):432.

26. Saito H, Yoshimura H, Tanaka K, Kimura H, Watanabe K, Tsubokura M, et al. Predicting CKD progression using time-series clustering and light gradient boosting machines. Scientific reports. 2024;14(1):1723.

27. Wang K, Liu Q, Mo S, Zheng K, Li X, Li J, et al. A decision tree model to help treatment decision-making for severe spontaneous intracerebral hemorrhage. International journal of surgery (London, England). 2024;110(2):788–98.

28. Wang Q, Qiao W, Zhang H, Liu B, Li J, Zang C, et al. Nomogram established on account of Lasso-Cox regression for predicting recurrence in patients with early-stage hepatocellular carcinoma. Frontiers in immunology. 2022;13:1019638.

29. Qiu X, Mao Q, Tang Y, Wang L, Chawla R, Pliner HA, et al. Reversed graph embedding resolves complex single-cell trajectories. Nature methods. 2017;14(10):979–82.

30. Jin S, Plikus MV, Nie Q. CellChat for systematic analysis of cell-cell communication from single-cell transcriptomics. Nature protocols. 2025;20(1):180–219.

31. Xie J, Li X, Zhang L, Liu C, Leung JW, Liu P, et al. Genistein-3’-sodium sulfonate ameliorates cerebral ischemia injuries by blocking neuroinflammation through the α7nAChR-JAK2/STAT3 signaling pathway in rats. Phytomedicine. 2021;93:153745.

32. Hilkens N, Casolla B, Leung T, de Leeuw FJL. Stroke. Lancet. 2024;403(10446):2820–36.

33. Zhao Y, Zhang X, Chen X, Wei Y. Neuronal injuries in cerebral infarction and ischemic stroke: From mechanisms to treatment (Review). International journal of molecular medicine. 2022;49(2).

34. Choi DW. Excitotoxicity: Still Hammering the Ischemic Brain in 2020. Frontiers in neuroscience. 2020;14:579953.

35. Qin C, Yang S, Chu YH, Zhang H, Pang XW, Chen L, et al. Signaling pathways involved in ischemic stroke: molecular mechanisms and therapeutic interventions. Signal transduction and targeted therapy. 2022;7(1):215.

36. Salaudeen MA, Bello N, Danraka RN, Ammani ML. Understanding the Pathophysiology of Ischemic Stroke: The Basis of Current Therapies and Opportunity for New Ones. Biomolecules. 2024;14(3).

37. Su Z, Ye Y, Shen C, Qiu S, Sun Y, Hu S, et al. Pathophysiology of Ischemic Stroke: Noncoding RNA Role in Oxidative Stress. Oxidative medicine and cellular longevity. 2022;2022:5815843.

38. Santos GSP, Magno LAV, Romano-Silva MA, Mintz A, Birbrair A. Pericyte Plasticity in the Brain. Neuroscience bulletin. 2019;35(3):551–60.

39. Abdullahi W, Tripathi D, Ronaldson PT. Blood-brain barrier dysfunction in ischemic stroke: targeting tight junctions and transporters for vascular protection. American journal of physiology Cell physiology. 2018;315(3):C343–c56.

40. Okada T, Suzuki H, Travis ZD, Zhang JH. The Stroke-Induced Blood-Brain Barrier Disruption: Current Progress of Inspection Technique, Mechanism, and Therapeutic Target. Current neuropharmacology. 2020;18(12):1187–212.

41. Jurcau A, Simion A. Neuroinflammation in Cerebral Ischemia and Ischemia/Reperfusion Injuries: From Pathophysiology to Therapeutic Strategies. International journal of molecular sciences. 2021;23(1).

42. Cao Y, Yue X, Jia M, Wang J. Neuroinflammation and anti-inflammatory therapy for ischemic stroke. Heliyon. 2023;9(7):e17986.

43. Carbone F, Teixeira PC, Braunersreuther V, Mach F, Vuilleumier N, Montecucco F. Pathophysiology and Treatments of Oxidative Injury in Ischemic Stroke: Focus on the Phagocytic NADPH Oxidase 2. Antioxidants & redox signaling. 2015;23(5):460–89.

44. Bradley D, Blaszczak A, Yin Z, Liu J, Joseph JJ, Wright V, et al. Clusterin Impairs Hepatic Insulin Sensitivity and Adipocyte Clusterin Associates With Cardiometabolic Risk. Diabetes care. 2019;42(3):466–75.

45. Du X, Chen Z, Shui W. Clusterin: structure, function and roles in disease. International journal of medical sciences. 2025;22(4):887–96.

46. Kwon HS, Kim TB, Lee YS, Jeong SH, Bae YJ, Moon KA, et al. Clusterin expression level correlates with increased oxidative stress in asthmatics. Annals of allergy, asthma & immunology : official publication of the American College of Allergy, Asthma, & Immunology. 2014;112(3):217–21.

47. Du J, Liu L, Yu B, Hao F, Liu G, Jing H, et al. Clusterin (apolipoprotein J) facilitates NF-κB and Bax degradation and prevents I/R injury in heart transplantation. International journal of clinical and experimental pathology. 2018;11(3):1186–96.

48. Pasten C, Herrera-Luna Y, Lozano M, Rocco J, Alvarado C, Liberona J, et al. Glutathione S-Transferase and Clusterin, New Players in the Ischemic Preconditioning Renal Protection in a Murine Model of Ischemia and Reperfusion. Cellular physiology and biochemistry : international journal of experimental cellular physiology, biochemistry, and pharmacology. 2021;55(5):635–50.

49. Iłżecka J, Iłżecki M, Grabarska A, Dave S, Feldo M, Zubilewicz T. Clusterin as a potential marker of brain ischemia-reperfusion injury in patients undergoing carotid endarterectomy. Upsala journal of medical sciences. 2019;124(3):193–8.

50. Martens GA, Geßner C, Osterhof C, Hankeln T, Burmester T. Transcriptomes of Clusterin-and S100B-transfected neuronal cells elucidate protective mechanisms against hypoxia and oxidative stress in the hooded seal (Cystophora cristata) brain. BMC neuroscience. 2022;23(1):59.

51. Foster EM, Dangla-Valls A, Lovestone S, Ribe EM, Buckley NJ. Clusterin in Alzheimer’s Disease: Mechanisms, Genetics, and Lessons From Other Pathologies. Frontiers in neuroscience. 2019;13:164.

52. Miguez-Cabello F, Wang XT, Yan Y, Brake N, Alexander RPD, Perozzo AM, et al. GluA2-containing AMPA receptors form a continuum of Ca(2+)-permeable channels. Nature. 2025;641(8062):537–44.

53. Butler-Ryan R, Wood IC. The functions of repressor element 1-silencing transcription factor in models of epileptogenesis and post-ischemia. Metabolic brain disease. 2021;36(6):1135–50.

54. Liu Z, Chen X, Gao Y, Sun S, Yang L, Yang Q, et al. Involvement of GluR2 up-regulation in neuroprotection by electroacupuncture pretreatment via cannabinoid CB1 receptor in mice. Scientific reports. 2015;5:9490.

55. Guo Y, Cai Y, Zhang X. Icariin ameliorates the cognitive function in an epilepsy neonatal rat model by blocking the GluR2/ERK I/II pathway. Folia neuropathologica. 2020;58(3):245–52.

